# How important are concurrent vehicle control groups in (sub)chronic non-human primate toxicity studies conducted in pharmaceutical development? An opportunity to reduce animal numbers

**DOI:** 10.1101/2023.02.15.528719

**Authors:** Lars Mecklenburg, Sarah Lenz, Georg Hempel

## Abstract

Safety assessment of human pharmaceuticals demands extensive animal experiments before a compound can be tested in patients or released on the market. Such experiments typically include concurrent vehicle control groups. Reconsidering the need for concurrent controls could support the strive to reduce the use of animals for scientific purposes. We reviewed reports from 20 (sub)chronic toxicity studies that were conducted in non-human primates (NHP) to characterize hazards of novel human pharmaceuticals. Firstly, we determined the toxicological endpoints that were identified to characterize the hazard. Secondly, we evaluated if the hazard could have been identified without reference to the concurrent controls. Thirdly, we employed an alternative statistical method to test for any significant change related to dose level or time. We found that toxicologically relevant hazards were identifiable without reference to concurrent controls, because individual measurements could be compared with pre-dosing values or because individual measurements could be compared to historical reference data. Effects that could not be evaluated without reference to concurrent controls were clinical observations and organ weights for which appropriate historical reference data was not available, or immune responses that could not be compared to pre-dosing measurements because their magnitude would change over time. Our investigation indicates that concurrent control groups in (sub)chronic NHP toxicity studies are of limited relevance for reaching the study objective. Under certain conditions, regulatory (sub)chronic NHP toxicity studies represent a good starting point to implement virtual control groups rather than concurrent control groups in nonclinical safety testing.

## Introduction

Safety assessment of new pharmaceuticals is highly regulated, demanding that extensive animal experiments are conducted before a pharmaceutical can be tested in patients or released on the market [1]. In Europe for example, Directive 2001/83/EG requires that the application to obtain a marketing authorization for a medicinal product be accompanied by toxicological and pharmacological tests, which are widely referred to as “nonclinical safety studies”. Guidance on such nonclinical safety studies is given by the International Council for Harmonization of Technical Requirements for Pharmaceuticals for Human Use (ICH) and by the Organization for Economic Co-operation and Development (OECD). Nonclinical safety studies aim to characterize the hazard associated with new pharmaceuticals and to determine a No Adverse Observed Effect Level (NOAEL) which functions as a reference for determining an acceptable exposure range for humans, e.g. patients in a clinical trial.

Societies across the globe strive towards reducing the use of animals for scientific purposes [2–6]. For example, the European Medicines Agency (EMA) committed to the application of replacement, reduction and refinement of animal testing as detailed in Directive 2010/63/EU [7], and in the United States of America, congress recently passed the FDA Modernization Act 2021, which allows for the use of New Approach Methodologies to evaluate the safety of drugs [8,9]. Replacement, reduction and refinement of animal testing are widely known as the “3Rs”. This principle was originally described in 1959 by Russel and Burch [10], who published considerations on how to promote “humane” behavior towards animals in the laboratory: “*Replacement means the substitution for conscious living higher animals of insentient material. Reduction means reduction in the numbers of animals used to obtain information of a given amount and precision. Refinement means any decrease in the incidence or severity of inhumane procedures applied to those animals which still have to be used*.”

With the aim to reduce the number of animals used in regulatory toxicity studies, the concept of virtual control groups has been introduced into nonclinical safety assessment [11]. Virtual control groups are supposed to replace concurrent control groups, which represent up to 25% of the animals in a regulatory toxicology study [12]. These animals are typically administered the vehicle, i.e. an inert medium that is used as a solvent or diluent in which the medicinally active agent is formulated. Virtual control groups are constructed from historical control data. Building adequate virtual control groups, however, is challenging and bears the risk that toxicological study outcomes are influenced, if covariates are not sufficiently controlled [13].

In consideration of the desire to replace concurrent controls with virtual controls, we aimed to evaluate the relevance of concurrent control groups in today’s pharmaceutical safety assessment. We specifically focused on those nonclinical safety studies that are conducted after the first human patients have been exposed, i.e. toxicity studies with subchronic (13 weeks) or chronic (up to 52 weeks) test article exposure. When those studies are performed, relevant information about the maximum recommended starting dose in human clinical trials and information about the pharmaceutical vehicle has already been collected, and the risk for misinterpreting vehicle-related effects is low. We also specifically focused on nonclinical studies that are conducted in non-human primates (NHP), since the use of NHP is of the greatest concern to the public [14]. We wanted to understand the relevance that concurrent control groups play in the determination of test article-related adverse effects and if omitting concurrent controls would still allow to reach the objective of these studies. We reviewed reports from 20 regulatory toxicity studies with the following approach: Firstly, we determined the toxicological endpoints that were identified in these studies that characterize the hazard associated with the pharmaceutical. Secondly, we evaluated if the hazard could have been identified without reference to the control group. Thirdly, we employed an alternative statistical method to objectively test for any significant change related to dose level or time.

## Materials and Methods

### Study reports for review

For the purpose of this investigation, final reports from 20 repeat dose toxicity studies of novel pharmaceuticals were selected. The selection process included all regulatory (Good Laboratory Practice-compliant) repeat dose toxicity studies with at least 90-day duration that were conducted at the test facility (Labcorp Early Development Services GmbH, Muenster, Germany) in the 3 years prior to the review. Studies with non-systemic routes of administration (e.g. intrathecal dosing) were excluded.

All studies were conducted in compliance with Directive 2010/63/EU, had undergone an ethical review and had been approved by the respective regional authority (Landesamt für Natur, Umwelt und Verbraucherschutz) based on national law in Germany. The test facility is accredited by AAALAC since 2007.

The studies were characterized as follows:

○ 19 studies were conducted in cynomolgus monkeys (*Macaca fascicularis*), one study was conducted in marmosets (*Callithrix jacchus*)
○ All studies included repeated dosing via one of the following routes of administration: oral (gavage), intravenous (injection or infusion), or subcutaneous
○ All studies included multiple dose groups (between 2 and 4) and 1 vehicle control group
○ Group sizes were between 3 and 6 animals per group and sex
○ The 20 studies included 349 male and 351 female animals
○ Several studies included a recovery phase in which a cohort of animals was left untreated for some time to evaluate reversibility of effects

A tabular overview of all 20 studies is provided in **Table 1**, including the number of animals and groups, route of administration, and pharmaceutical class of the test article.

**Table 1:** Overview of the 20 studies included in this subjective retrospective analysis.

| Study ID | Species | Design (M/F) | Duration (weeks) | RoA | Pharmaceutical class |
| --- | --- | --- | --- | --- | --- |
| A | Cynom. | V: 6/6<br>L: 4/4<br>M: 4/4<br>H: 6/6 | 26 | sc | mAb |
| B | Cynom. | V: 3/3<br>L: 3/3<br>H: 3/3 | 26 | iv | mAb |
| C | Cynom. | V: 6/6<br>L: 4/4<br>M: 4/4<br>H: 6/6 | 26 | sc | mAb |
| D | Cynom. | V: 6/6<br>L: 4/4<br>M: 4/4<br>H: 6/6 | 26 | sc | mAb |
| E | Cynom. | V: 3/3<br>L: 3/3<br>H: 3/3 | 26 | iv | mAb |
| F | Cynom. | V: 6/6<br>L: 4/4<br>M: 4/4<br>H: 6/6 | 12 | og | small molecule |
| G | Cynom. | V: 5/5<br>L: 3/3<br>M: 3/3<br>H: 5/5 | 13 | sc | Oligonucleotide |
| H | Cynom. | V: 6/6<br>L: 4/4<br>LM: 4/4<br>MH: 4/4<br>H: 6/6 | 26 | sc | recombinant human peptide |
| I | Cynom. | V: 3/3<br>L: 3/3<br>M: 3/3<br>H: 3/3 | 13 | og | small molecule |
| J | Cynom. | V: 6/6<br>L: 4/4<br>M: 6/6<br>H: 6/6 | 27 | sc | mAb |
| K | Cynom. | V: 6/6<br>L: 4/4<br>M: 4/4<br>H: 6/6 | 39 | sc | ASO |
| L | Cynom. | V: 6/6<br>L: 4/4<br>M: 4/4<br>H: 6/6 | 39 | sc | ASO |
| M | Cynom. | V: 5/5<br>H: 6/8 | 26 | sc | mAb |
| N | Cynom. | V: 5/5<br>L: 3/3<br>M: 3/3<br>H: 5/5 | 13 | og | small molecule |
| O | Cynom. | V: 5/5<br>L: 3/3<br>M: 3/3<br>H: 5/5 | 4 | og | small molecule |
| P | Cynom. | V: 6/6<br>L: 4/4<br>M: 4/4<br>H: 7/7 | 26 | sc | ASO |
| Q | Marmoset | V: 6/6<br>L: 4/4<br>M: 4/4<br>H: 6/6 | 13 | og | small molecule |
| R | Cynom. | V: 6/6<br>L: 6/6<br>H: 6/6 | 13 | iv | mAb |
| S | Cynom. | V(sc): 5/5<br>L(sc): 4/4<br>H(sc): 6/6<br>H(iv): 4/4 | 13 | sc/iv | mAb fragment |
| T | Cynom. | V: 6/6<br>L: 6/6<br>H: 4/4 | 26 | sc | mAb |

### Descriptive analysis of study reports

Study reports were blinded for the test article and sponsor and were reviewed for the following parameters: Clinical Observations, Body Weight, Ophthalmic Examination, Body Temperature, Electrocardiography, Blood Pressure, Urinalysis, Hematology, Coagulation biomarkers, Clinical Chemistry, Immune cell phenotyping, Cytokines/ Chemokines, T Cell Dependent Antibody Response, Organ Weights, Macroscopic Findings, Microscopic Findings. Toxicological endpoints that were specifically called out by the study director were subsequently allocated to one of the following categories:

○ Test article related, adverse, i.e. they were considered for determining the NOAEL
○ Test article related, non-adverse, i.e. they were not considered for determining the NOAEL
○ Vehicle- or procedure related
○ Incidental with no toxicological relevance

### Analysis without reference to the control group

One of the authors (SL) conducted a retrospective analysis of all parameters that were specifically called out by the study director. The aim of the retrospective analysis was to understand why they had been called out and whether the effect was also detectable without reference to the concurrent control group. Subsequently, concordance with the original interpretation was evaluated and differences or challenges in the interpretation without concurrent control groups were highlighted. The analysis followed general recommendations on determining adversity and the NOAEL in nonclinical safety studies [15] and was conducted with particular emphasis on the following aspects:

○ Timing of effect and development over time
○ Magnitude of the effect and dependence on dose level
○ Number of individual animals demonstrating the effect
○ Comparison of measurements under dosing with those obtained before dosing
○ Comparison of measurements under dosing with reference data obtained from previously used control animals

### Statistical analysis by mixed-design ANOVA

All studies had used the fixed-effect analysis of variance (ANOVA) and subsequent Dunnett’s test to statistically analyze differences between dosed animals and control animals at a given time point [16]. Endpoints for which a statistically significant difference was observed were highlighted.

Selected endpoints were subjected to an alternative statistical analysis after omission of data from the concurrent control group. Rather than using a fixed-effect ANOVA, a mixed-design ANOVA was used. In a mixed-design ANOVA model, one factor (a fixed effect factor) is a between-subject variable and the other (a random effect factor) is a within-subject variable [17]. The mixed design ANOVA investigates an effect by time, an effect by group (dose) and the interaction of these two effects. The mixed design ANOVA requires that residuals are normally distributed, that variance is homogenous and that measurements show sphericity at more than two time points. All requirements were proven or assumed.

The statistical analysis was conducted using an established software system (IBM SPSS Statistics). If significant differences were observed between the main effect time or group, a post-hoc analysis was conducted.

## Results

### Analysis of study data after omission of the concurrent control group

Out of 20 studies, two studies (studies J, N) did not reveal any relevant observation that was specifically called out in the study report. The findings that were called out in the study reports of the remaining 18 studies are listed in **Table 2**.

**Table 2:** Overview of results per study; the table presents all findings that were called out by the study director; findings are allocated to four different categories; NOAEL describes the dose level that was selected as NOAEL by the study director. For each finding it is indicated whether or not concordance occurred between the original interpretation and the subjective interpretation after omission of the control group.

| Study ID | Test article-related effect, adverse | Test article-related effect, non-adverse | Vehicle/procedure-related effect | Incidental | NOAEL |
| --- | --- | --- | --- | --- | --- |
| A | - | Body temperature↑§<br>Serum triglyceride↑§ | - | - | H |
| B | - | Clinical obs (skin)§<br>TDAR↓ | - | - | H |
| C | - | Clinical obs (injection site)§<br>Serum IL-6↑§<br>Mic (injection site)# | - | - | H |
| D | - | Soft feces§<br>Blood eosinophils↓§ | - | - | H |
| E | - | Serum globulin↑§ | - | - | H |
| F | Body weight↓§<br>Mic(testis)#<br>Mac(testis)# | Organ weight(liver)↑<br>Organ weight(spleen)↑ | Vomiting<br>Blood pressure↓ | - | M |
| G | Platelets↓§,# | Mic(lymph node)#<br>Mac(lymph node)# | - | - | M |
| H | - | Mic(injection site)#<br>Serum AP↑§<br>TDAR↓ | - | Clinical Obs(skin)<br>Patellar reflex↓ | H |
| I | - | Salivation§<br>Soft feces§<br>Clinical Obs(skin)§<br>Serum cholesteroles↓§<br>Organ weight(liver)↑ | Blood RBC↓§ | - | H |
| J | - | - | - | - | H |
| K | - | Mic(liver)#<br>Mic(kidney)#<br>Mic(lymph node)# | - | - | H |
| L | - | Serum globulin↓§<br>Mic(liver)#<br>Mic(kidney)#<br>Mic(lymph node)#<br>Mic(injection site)# | - | - | H |
| M | - | Serum IgG↓§<br>Serum albumin↓§ | - | - | H |
|  |  | Serum colesterole↑§<br>TDAR↓ |  |  |  |
| N | - | - | - | - | H |
| O | Mic(testis)#<br>Mac(testis)# | Blood RBC↓§ | - | - | H (females)<br>L (males) |
| P | - | Serum ALT, AP, GLDH↑§<br>Mic(liver)#<br>Mic(kidney)#<br>Mic(lymph node)#<br>Mic(injection site)#<br>Organ weight(liver)↑ | - | Soft feces§ | H |
| Q | Serum GLDH, AST, ALT, GGT↑§<br>Mic(liver)# | Vomiting§ | Body weight↓§<br>Serum albumin↓§ | - | M |
| R | - | Blood CD20+ lymphocytes↓§<br>Mic(spleen)#<br>Mic(lymph node)# | - | - | H |
| S | - | Serum IgG↓§<br>Serum albumin↓§<br>Serum Ca↓§<br>TDAR↓ | - | - | H |
| T | - | Mic(injection site)# | - | - | H |

#, concordant with reference to historical data; §, concordant with reference to predose value; grey, not concordant; ALT, alanine aminotransferase; AST, aspartate aminotransferase; GGT, gamma glutamyltransferase; GLDH, Glutamate dehydrogenase; H, High dose level; L, Low dose level; Mac, macroscopic observation; Mic, microscopic observation; TDAR, T Cell Dependent Antigen Response.

### Test article related adverse findings

Test article related adverse findings occurred in 4 studies (studies F, G, O, Q) and consisted of reduced body weight, microscopic and macroscopic observations and alterations in clinical pathology concerning platelets or serum liver enzyme activities. All of these test article-related adverse findings were used to determine a NOAEL for the study. We investigated, if those test article related adverse findings were detectable without reference to the concurrent control group:

In study F, body weight was reduced at the end of the dosing phase in three female animals from the high dose group when compared to control. It was also detectable without the vehicle control group by comparison to pre-dosing values.

In the same study, male animals from the high dose group showed macroscopic and microscopic observations in the testis. These changes could also be determined without the control group, since they exceeded the known background pathology represented in a historical reference data set.

In study G, two animals from the high dose group showed reduced platelet counts in comparison to the control group. With omission of the control group, the reduced platelet count was still identifiable by comparison to pre-dose values and by comparison with a historical reference data set.

In study O, animals from the mid dose and the high dose level showed microscopic and macroscopic observations in the testis, with higher severity in the high dose compared to the mid dose group. With omission of the control group, these observations were still detectable, since they exceeded the known background pathology represented in a historical reference data set.

In study Q, elevated serum levels of glutamate dehydrogenase, aspartate aminotransferase, alanine aminotransferase, and gamma-glutamyltransferase were observed at the end of dosing in high dose animals when compared to controls. Without the control group, this change was also apparent by comparing to pre-dosing values and to historical reference data. In addition, the above clinical pathology changes correlated with microscopic observations, i.e. hypertrophy and hyperplasia of intrahepatic bile ducts.

### Test article related non-adverse findings

Test article-related non-adverse findings occurred in 18 studies. Most of these findings were detectable without reference to the control group, with the exception of organ weight changes and changes in T Cell Dependent Antigen Response (TDAR).

In study F, the mean absolute liver weight in the high dose group was increased compared to the control group. In the same study, a difference in the mean absolute spleen weight was found between the high dose group and controls. Since no microscopic correlate was found for the increased organ weights, they were not considered adverse. Without the concurrent control, organ weights can only we compared to historical reference data, which is not always available. A difference between control and dose group was also found for the relative liver weight (to body weight) in two other studies (studies I, P) with the same limitations after omission of the control group as mentioned above.

The TDAR assay is a measure of immune function that is dependent upon the effectiveness of multiple immune processes, including antigen uptake and presentation, T cell help, B cell activation, and antibody production. Immunoglobulin (Ig) M and G levels are measured before and after immunization with a standard antigen such as Keyhole Limpet Hemocyanin. In study B, a reduced post-immunization IgG level was found. This observation was made in both groups administered the test article, but not in the control group. Since this effect was expected, given the pharmacological mode of action of the test article, it was not considered adverse. Without a control group, the magnitude of a reduced post-immunization response cannot be determined, particularly on an individual animal level, given the high inter-individual variability in this parameter (**Fig 1**). Nevertheless, reduced IgG levels were still detectable without a concurrent control, since they remained below 50,000 at all times, indicating a clear deviation from reference data. Similar observations were made in the TDAR of three other studies (studies H,M,S). In all cases, reduced post-immunization antibody responses were expected given the pharmacologic mode of action of the respective test article.

**Figure 1:**
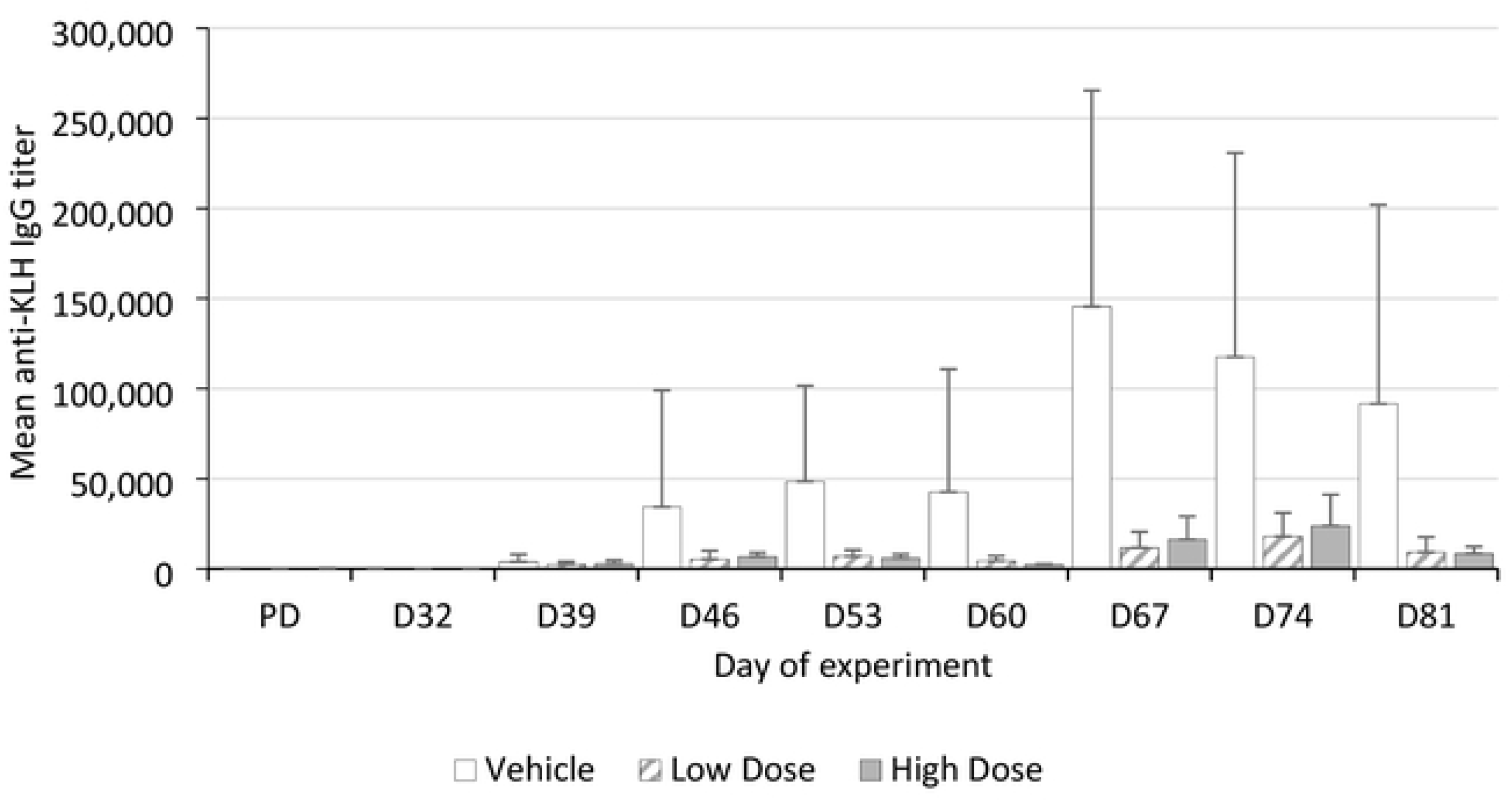
Mean anti-KLH IgG titer from a 26-week intravenous toxicity study in cynomolgus monkeys. Animals (n=6 per group) were allocated to 3 groups (vehicle control, low dose test article, high dose test article) and were immunized on days 32 and 60 of the dosing phase. Error bars = standard deviation.

### Vehicle- or procedure related findings

Vehicle- or procedure related findings occurred in 3 studies (studies F, I, Q). They consisted of vomiting, reduced blood pressure, reduced red blood cell count, reduced body weight and reduced serum albumin concentration.

Vomiting was observed repeatedly throughout study F. It was observed in 3 males from the control group, in one male from the mid dose group and in four males as well as three females from the high dose group. Since animals from the control group were similarly affected, vomiting was attributed to the vehicle. Without the concurrent control group, vomiting could not be interpreted since no reference data was available for this clinical endpoint.

A slight reduction in systolic, diastolic, and mean arterial blood pressure occurred in males of study F. The slightly reduced blood pressure was ascribed to the high frequency of this measurement taken throughout the study and was interpreted as an adaptation of the animals to the procedure. Consequently, the reduction in blood pressure was attributed to the procedure of the study. Without reference to the concurrent control group, the reduction in blood pressure would have been ascribed to the test article.

A reduced red blood cell mass, hematocrit and blood hemoglobin concentration was observed in study I. This finding was also observed in controls and therefore was considered related to the repeated blood collection for toxicokinetic evaluation. Without a concurrent control, however, procedure-related changes cannot be differentiated from effects of the test article.

In study Q, a body weight loss was noted in nearly all animals of all groups, including controls. After approximately 4 weeks, a stabilization of body weights was recorded, and most animals gained body weight again in the following weeks. This finding was considered related to stress due to the very frequent handling procedures occurring at least twice daily. Without reference to the concurrent control group, the body weight loss would have been attributed to the test article.

A reduced albumin concentration in serum was observed in study Q. This was recoded in all groups including controls. Since animals from the control group were similarly affected as were dosed animals, these observations had been ascribed to the vehicle. Without a control group, vehicle-associated toxicities are not detectable and these findings would have been ascribed to the test article.

### Incidental findings

Incidental findings occurred in two studies (study H, P). They consisted of clinical observations in the skin, a reduced patellar reflex and the occurrence of soft feces.

Red skin discoloration or scaling were observed at the trunk and the inguinal region in study H. These findings were observed in four animals of the control group, in four animals of the low dose group, in two animals of the mid dose group, and in three animals of the high dose group. They were not accompanied by itching and did not impact the overall health of the animals. Microscopic analysis of affected skin was not conducted. Since the clinical findings occurred in vehicle control animals and in test article dosed animals, the skin findings were not considered test article related. Without reference to the control group, these clinical observations could not be interpreted since no reference data was available for this clinical endpoint.

A transiently non-functional patellar reflex was observed in study H. It occurred in four animals from the control group, in one animal of the low dose group, in two animals of the mid dose group and in two animals of the high dose group. Since animals from the control group were similarly affected, this finding was not considered test article related. Without a control group, these clinical observations could not be interpreted since no reference data was available for this clinical endpoint.

Occasional occurrence of soft feces was observed in study P. This observation was made in six control animals, in six animals of the low dose group, and in four animals of the mid dose group, but in no high dose animal. Since animals from the control group were similarly affected, this observation was considered incidental. Without a control group, this observation could not be interpreted since no reference data was available for this clinical endpoint.

### Statistical analysis after omission of control groups

It is standard in pivotal repeat dose toxicity studies to compare measurements between dosed animals and control animals. The statistical analysis typically employed is a fixed-effect analysis of variance (ANOVA) followed by Dunnett’s test [16]. If control groups are omitted, application of a fixed-effect ANOVA is no longer feasible. Nevertheless, other statistical methods can be applied to compare longitudinal measurements, i.e. measurements that are repeatedly examined over time. The mixed-design ANOVA investigates an effect by time, an effect by group (dose) and the interaction of these two effects.

In the 20 studies that were included in this investigation, 22 observations had shown a statistically significant (p>0.05) difference between dose groups and the control group using the fixed-effect ANOVA/Dunnett’s test. Those 22 observations were mostly in clinical chemistry measurements, one was a body temperature measurement and one referred to the phenotyping of immune cells (**Table 3**). We examined all measurements using the mixed-design ANOVA approach after omitting data from the control group.

**Table 3:** Summary of statistical analysis. In total, 22 statistically significant results were obtained with the fixed-effect ANOVA/Dunnett’s test. Result from mixed-design ANOVA (conducted after omission of the control group) are listed on the right.

| Number of Results | Study | Category | Parameter | Sex | Significant Interaction Effect | Significant Effect of Time | Significant Effect of Group |
| --- | --- | --- | --- | --- | --- | --- | --- |
| 1. | Study A | Body temperature | T | F | X | - | - |
| 2. | Study A | Clinical Chemistry | TRIG | M | O | X | X |
| 3. | Study E | Clinical Chemistry | GLOB | M | O | X | X |
| 4. | Study E | Clinical Chemistry | A:G | M | O | X | X |
| 5. | Study H | Clinical Chemistry | ALP | F | O | O | O |
| 6. | Study L | Clinical Chemistry | GLOB | F | O | X | O |
| 7. | Study L | Clinical Chemistry | TP | F | O | X | O |
| 8. | Study M | Clinical Chemistry | IgG | M | O | X | O |
| 9. | Study M | Clinical Chemistry | TP | M | O | X | O |
| 10. | Study M | Clinical Chemistry | GLOB | M | O | X | O |
| 11. | Study M | Clinical Chemistry | ALB | M | O | X | O |
| 12. | Study M | Clinical Chemistry | CHOL | M | O | X | O |
| 13. | Study P | Clinical Chemistry | ALT | F | O | O | O |
| 14. | Study P | Clinical Chemistry | GLDH | F | O | O | X |
| 15. | Study Q | Clinical Chemistry | AST | F | O | X | O |
| 16. | Study Q | Clinical Chemistry | GGT | F | O | O | O |
| 17. | Study R | Immunophenotyping | CD20+ B-cells | F | X | - | - |
| 18. | Study S | Clinical Chemistry | IgG | F | O | X | O |
| 19. | Study S | Clinical Chemistry | CA | F | O | O | O |
| 20. | Study S | Clinical Chemistry | TP | F | O | X | O |
| 21. | Study S | Clinical Chemistry | ALB | F | O | X | O |
| 22. | Study S | Clinical Chemistry | A:G | F | X | - | - |

- = not analysed; () = obsolete due to interaction; A:G, albumin to globulin ratio; ALP, alkaline phosphatase; ALT, alanine aminotransferase; CA, calcium; CHOL, cholesterol; GGT, gamma-glutamyltransferase; GLOB, globulin; O = not significant; T, temperature; TP, total protein; TRIG, triglyceride; X = significant.

The mixed-design ANOVA revealed a concordant significant effect for 18 of these 22 observations. For three observations, a significant interaction effect was demonstrated, i.e. measurements were dependent on time and dose. For another 15 observations, either an effect of time or an effect of dose was demonstrated (**Table 3**). No concordant statistical difference was found for the following 4 observations:

A significant elevation in alkaline phosphatase serum activity was described in study H. Mean alkaline phosphatase activity was significantly (P≤0.01) increased in animals from the high dose group (1315 U/L) when compared to controls (810 U/L). The elevation in alkaline phosphatase serum activity was also apparent when values at the end of the dosing phase were compared to mean pre-dose values (927 U/L). While the fixed-effect ANOVA/Dunnett’s test had revealed a statistical difference between high dose group and control at a single time point (day 83 of dosing), the mixed-design ANOVA did not reveal a statistical effect of time or group.

A significant elevation in alanine aminotransferase serum activity was observed in study P. Mean alanine aminotransferase activity was significantly (P≤0.01) increased in animals from the high dose group (96 U/L) when compared to controls (45 U/L). The elevation, however, was still below the upper normal reference limit for this parameter (102.23 U/L). The elevation in alkaline phosphatase activity was also apparent when dosing values were compared to pre-dose values (52 U/L). While the fixed-effect ANOVA/Dunnett’s tests revealed a statistical difference between high dose group and control at a single time point (day 55 of dosing), the mixed-design ANOVA did not reveal a statistical effect of time or group.

A significant elevation in gamma-glutamyltransferase serum activity was observed in study Q. Mean gamma-glutamyltransferase activity was significantly (P≤0.05) increased in animals from the high dose group (155 U/L) when compared to controls (12 U/L). The elevation in gamma-glutamyltransferase serum activity was also apparent when dosing values were compared to mean pre-dose values (8 U/L). While the fixed-effect ANOVA/Dunnett’s test revealed a statistical difference between high dose group and control, the mixed-design ANOVA did not reveal a statistical effect of time or group, likely attributed to a very high inter-individual variability.

A significant decrease in serum calcium levels was observed in study S. Mean calcium levels were significantly (P≤0.05) decreased in animals from the mid dose group (2.28 mmol/L) and the high dose group (2.10 mmol/L) when compared to controls (2.54 mmol/L). The decrease in serum calcium was also apparent when values on dosing day 86 were compared to pre-dose values (2.55 mmol/L in both mid and high dose). While the fixed-effect ANOVA/Dunnett’s tests revealed a statistical difference between high dose group and control, the mixed-design ANOVA did not reveal a statistical effect of time or group, likely caused by a high inter-individual variability at the pre-dosing time point.

## Discussion

With this investigation we explored the relevance of concurrent control groups in regulatory NHP (sub)chronic toxicity studies for novel pharmaceuticals. For our investigation we reviewed 20 finalized study reports. This number was not based on any statistical calculation but was chosen arbitrarily based on the available capacity. The 20 studies cover a broad range of test articles (including small molecule drugs and biotechnology-derived drugs) and represent typical studies that are conducted for regulatory purposes in pharmaceutical development (**Table 1**). We decided to conduct a non-blinded retrospective analysis, since toxicological endpoints in such study types are extremely broad (ranging from subjective clinical observations to quantitative analytical measurements) and since there was no information available how interpretation for any of these endpoints would be affected by omission of the concurrent control group. With that in mind, our study is exploratory in nature and will require a follow up with prospective investigator-blinded investigations on the same matter.

Our investigation has shown that only 4 out of 20 studies (i.e. 20%) revealed dose-limiting toxicities. This reflects that for repeat dose toxicity studies of (sub)chronic duration, a large set of data on test article toxicity and toxicokinetics is already available, that allows for a robust selection of tolerable dose levels. It also suggests, that many of the modern, often biotechnology-derived test articles in pharmaceutical development are associated with a low risk for toxicity. In fact, two of the 20 studies (i.e. 10%) did not reveal any effect at all.

It is interesting and of high relevance that in all 4 studies where dose-limiting toxicities occurred, those could be defined without reference to the concurrent control group. This is important, because adverse dose-limiting toxicities are used to define the NOAEL which again is used to determine a safe dose range in clinical applications. Even if control groups had been omitted from these studies, the same NOAEL would have been defined.

Dose–limiting toxicities were seen as reduction in body weight, reduction in thrombocyte count and increased serum activities of liver enzymes. These are parameters that are repeatedly measured throughout an experiment and therefore allow a direct comparison between values taken before and after administration of the test article. Further dose-limiting toxicities were seen as macroscopic and microscopic findings. These endpoints are only measured once at the end of the experiment and therefore do not allow a pre- versus post-dose comparison. However, there is a vast set of historical reference data available for macroscopic and microscopic organ findings which allows interpretation of observations even in the absence of a concurrent control.

Moreover, our investigation has demonstrated that the majority of test article-related effects are non-adverse, i.e. they are not considered to represent a hazard for humans and therefore are not considered to define the NOAEL. Most of these non-adverse test article related findings were detectable without a concurrent control, because they are measured repeatedly throughout a study. A notable exception are organ weights that are only recorded once at the end of an experiment and for which historical reference data is not broadly available. Another exception is the TDAR, i.e. a method where specific serum IgG and IgM levels are measured after immunization with a T cell antigen. Since the individual response varies between individuals, a comparison to a concurrent control is mandatory to detect subtle differences. Nevertheless, large differences can be detected even without a concurrent control, if data is compared to a typical immune response from historical controls.

The typical concurrent control group in a regulatory repeat dose toxicity study for a new pharmaceutical is treated with the pharmaceutical vehicle, i.e. the same product but without the active pharmaceutical ingredient. Naturally, omission of the control group does not allow characterization of any vehicle-related effects. Consequently, omission of control groups (and replacing it with virtual controls) should only be considered if the vehicle is well characterized (or if virtual controls can be constructed from data associated with the same vehicle). Similarly, effects that are caused by the experimental procedure itself can no longer be differentiated from test article induced effects, if the concurrent control is omitted. We have seen such effects as a reduction of red blood cells following a frequent high volume blood collection or as a reduced blood pressure as animals got acquainted to the procedure and experienced less stress. However, those effects are typically well known and therefore rarely represent a challenge for the interpretation of study data.

A main reason to include concurrent controls in an experiment is the detection of incidental changes, i.e. effects that occur randomly for various reasons and that are not under the control of the researcher. We have seen such effects in two out of the 20 studies (i.e. 10%) as spontaneously occurring clinical observations in the skin, as non-functional patellar reflexes, or as spontaneous occurrence of soft feces. Without a concurrent control, it is impossible to differentiate such findings from test article-induced effects. This problem, however, could be addressed by generating large sets of historical reference data on virtually every endpoint that is collected within such study types. Currently, historical reference data is typically collected for macroscopic and microscopic observations and for clinical pathology parameters [11]. It would certainly be possible and advisable to generate similar data sets for clinical observations, for organ weights and generally for any other type of data that is collected.

Current practice based on OECD and ICH guidelines includes that measurements from repeat dose toxicity studies in non-rodents are statistically analyzed, despite the fact that group sizes are generally small and studies are typically underpowered to detect low magnitude effects. The method typically employed for this statistics is the fixed-effect analysis of variance (ANOVA) followed by pairwise comparisons using Dunnett’s test [16]. We propose that mixed-design ANOVA can be used for statistical analysis, if concurrent control groups are not available. Our investigation has demonstrated that out of 22 parameters with statistical difference in the fixed-effect ANOVA /Dunnett’s test, 82% could be confirmed using the mixed-design ANOVA. The 18% of non-concordant results are mainly associated with situations where a change occurs on a single occasion among several measurements and where the change is of low magnitude combined with a high inter-individual variability.

Recapitulating, is there potential to reduce animal numbers in (sub)chronic NHP regulatory toxicity studies by omitting concurrent controls (and potentially replacing them with virtual control groups)? Our investigation indicates that there is, because concurrent control groups are not typically needed to determine dose-limiting toxicities. While this conclusion should not be extrapolated to toxicity studies in other non-rodent species (e.g. dogs) or rodents, because the nature of test articles that are investigated in these species differs from the nature of test articles that are investigated in NHP [18], it should be understood as an indication that (sub)chronic NHP toxicity studies are a good starting point to implement virtual control groups in nonclinical safety testing. In the year 2019, a total of 1,219 cynomolgus macaques (*Macaca fascicularis*) and 32 marmoset monkeys (*Callithrix jacchus*) were used in regulatory repeat dose toxicity studies of at least 90-day duration within the 28 EU member states and Norway [19]. If only 20% of those animals were assigned to concurrent controls, their omission would reduce the need for 250 animals.

## Conclusion

Collectively, our investigation indicates that concurrent control groups in (sub)chronic NHP regulatory toxicity studies are of limited relevance for reaching the study objective. They are mainly needed to characterize effects on the immune response, effects on organ weights, vehicle- or procedure-related effects and incidental findings that cannot be controlled in an experiment. Yet, our investigation has shown that detection of test article-related adverse effects, i.e. those effects that are used to derive a safe dose range for clinical applications of a pharmaceutical, can still be detected without a concurrent control group. Therefore, regulatory (sub)chronic NHP toxicity studies represent a good starting point to implement virtual control groups – rather than concurrent control groups - in nonclinical safety testing.

## Notes

### Competing Interest Statement

The authors have declared no competing interest.

